# Anisotropy of plasmalemmal sterols and cell mating require StARkin domain proteins Ysp2 and Lam4

**DOI:** 10.1101/2021.09.30.462584

**Authors:** Neha Chauhan, Gregory D. Fairn

## Abstract

In the budding yeast S. cerevisiae Cdc42 is required for polarized growth and the formation of mating projections (shmoos). Negatively charged lipids including phosphatidylserine and phosphatidylinositol 4,5-bisphosphate support a positive feedback loop that recruits Cdc42 effectors and MAP kinase scaffolds, many of which contain polybasic patches that directly interact with the membrane. Here, using genetically encoded sterol sensor ALOD4 we find that ergosterol is accumulated in the cytosolic leaflet of buds and shmoos. The accumulation of ergosterol in the plasma membrane requires both Osh and Lam proteins however cells lacking Ysp2/Lam2 and Lam4 displayed a reversal in the polarity of ergosterol. The redistribution of ergosterol impairs the polarization of phosphatidylserine and phosphatidylinositol 4,5-bisphophate which further impacts shmoo formation, MAPK signaling and mating efficiency. Our observations demonstrate that the ability of Lam proteins to deliver ergosterol from the plasma membrane to the ER helps maintain a gradient of ergosterol which in turn supports robust cell polarity.

**Summary:** The sterol sensor ALOD4 is enriched at sites of polarized growth. Elimination of the Osh proteins solubilized the ALOD4 whereas elimination of Ysp2 and Lam4 reversed ALOD4 polarization. Cells lacking Ysp2 and Lam4 have defects in mating and MAP kinase signaling.

## Introduction

The anisotropic distribution of cell components is critical for cellular polarity, migration, polarized growth (Orlando and Guo, 2009) and in mating of budding yeast (Fairn et al., 2011; Garrenton et al., 2010; Slaughter et al., 2009). Small G-proteins of the Rho family are critical mediators of cell polarity (Hodge and Ridley, 2016) and in yeast Cdc42, the guanine exchange factor Cdc24 and the scaffold protein Bem1 are crucial regulators of polarized growth and mating (Kozubowski et al., 2008; Witte et al., 2017; Woods et al., 2015). The coordinated activation and amplification of Cdc42 at sites of bud emergence and mating projects facilitates actin polarization, focal exocytosis and polarized secretion and signal transduction (Kozubowski et al., 2008; Meca et al., 2019; Wedlich-Soldner et al., 2004). In addition to proteins, lipids within the plasma membrane (PM) are also important to support Cdc42 activation and its positive feedback loop. Anionic lipids including phosphatidylserine (PtdSer), phosphatidylinositol 4-phosphate (PI4P) and phosphatidylinositol 4,5-bisphosphate (PI4,5P_2_) have been documented to be enriched in the bud over the mother and at the tips of mating projects (Fairn et al., 2011; Garrenton et al., 2010). This assembly of negatively charged lipids helps support the localized recruitment of Cdc42 and other effectors.

In addition to anionic phospholipids and sphingolipids, ergosterol and other yeast sterols are enriched in the PM. Ergosterol, the most abundant yeast sterol, is crucial for maintaining fluidity, permeability, and lipid organization of the plasma membrane (PM). Yeast sterols are mainly synthesized *de novo* as yeast cannot import exogenous sterols under aerobic conditions. The rapid transport of ergosterol between cellular membranes is critical for maintaining membrane homeostasis and maintaining its high, ∼40 mol%, plasmalemmal content. Ergosterol is also enriched in the tips of mating projects where it supports pheromone signaling (Jin et al., 2008). Metabolic labeling of yeast treated with α-factor revealed a 30% increase in ergosterol levels (Bagnat and Simons, 2002) consistent with the notion that ergosterol is important for pheromone signaling, cell polarization and mating. How precisely ergosterol transport is regulated in vegetative yeast and those responding to mating factor is unclear. The transport of ergosterol and other lipids throughout the cell is mediated by both vesicular and non-vesicular pathways. Yeast has two protein families reported to be involved in non-vesicular intracellular sterol transport. The first are the cytosolic Osh proteins (Osh1-7) which collectively are essential for cell viability (Beh et al., 2001). The second set of sterol transport proteins are the six LAM proteins which contain a PH-gram domain, two sterol-binding StARkin domains and are tail anchored to the ER (Gatta et al., 2015).

Domain 4 of the Perfringolysin O (PFOD4) and Anthrolysin O (ALOD4) have been used to monitor cholesterol in the PM and other organelles in mammalian cells (Infante and Radhakrishnan, 2017; Maekawa and Fairn, 2015) and more recently in yeast cells (Encinar Del Dedo et al., 2021; Marek et al., 2020). These probes are complementary to the widely used stain filipin for monitoring the cellular distribution of sterols and have the advantage that they are suitable for live cell imaging. However, in contrast to commonly used lipid probes such as PH-PLCδ and Lact-C2 which bind in a 1:1 stoichiometry with PI4,5P_2_ and PtdSer, respectively, the D4 probes have a binding threshold typically >30 mol% cholesterol (Flanagan et al., 2009; Johnson et al., 2012). Here using a genetically encoded version of GFP-ALOD4, we find that yeast sterols in the cytosolic leaflet of the PM are highly accessible in the bud and the tips of mating projections. The StART-like domain proteins Ysp2 and Lam4 are required for the observed sterol anisotropy and loss of these proteins reduces the polarization of PtdSer and PI4,5P_2_, and attenuates MAP kinase signaling and cell mating.

## Results and Discussion

### ALOD4 as a biosensor for cytosolic leaflet sterols

Vegetatively growing yeast cells expressing GFP-ALOD4 were examined using spinning disc confocal microscopy. **Ergosterol was found to localize prominently in the** bud consistent with previous work (Encinar Del Dedo et al., 2021) (Fig. 1A and B). Treatment of cells with the squalene synthase inhibitor, zaragozic acid (ZA) resulted in a 38% reduction of sterols (Fig. 1C). Reduction in sterol content in the presence of ZA results in the relocalization of the GFP-ALOD4 sensor to the cytosol (Fig. 1D) consistent with the notion that ALOD4 requires a threshold of sterols to bind. This effect is reversible since the removal of ZA and continued growth restores the sensor to the PM. The data demonstrate that GFP-ALOD4 specifically binds sterols in the cytosolic leaflet of the PM.

**Figure 1.**
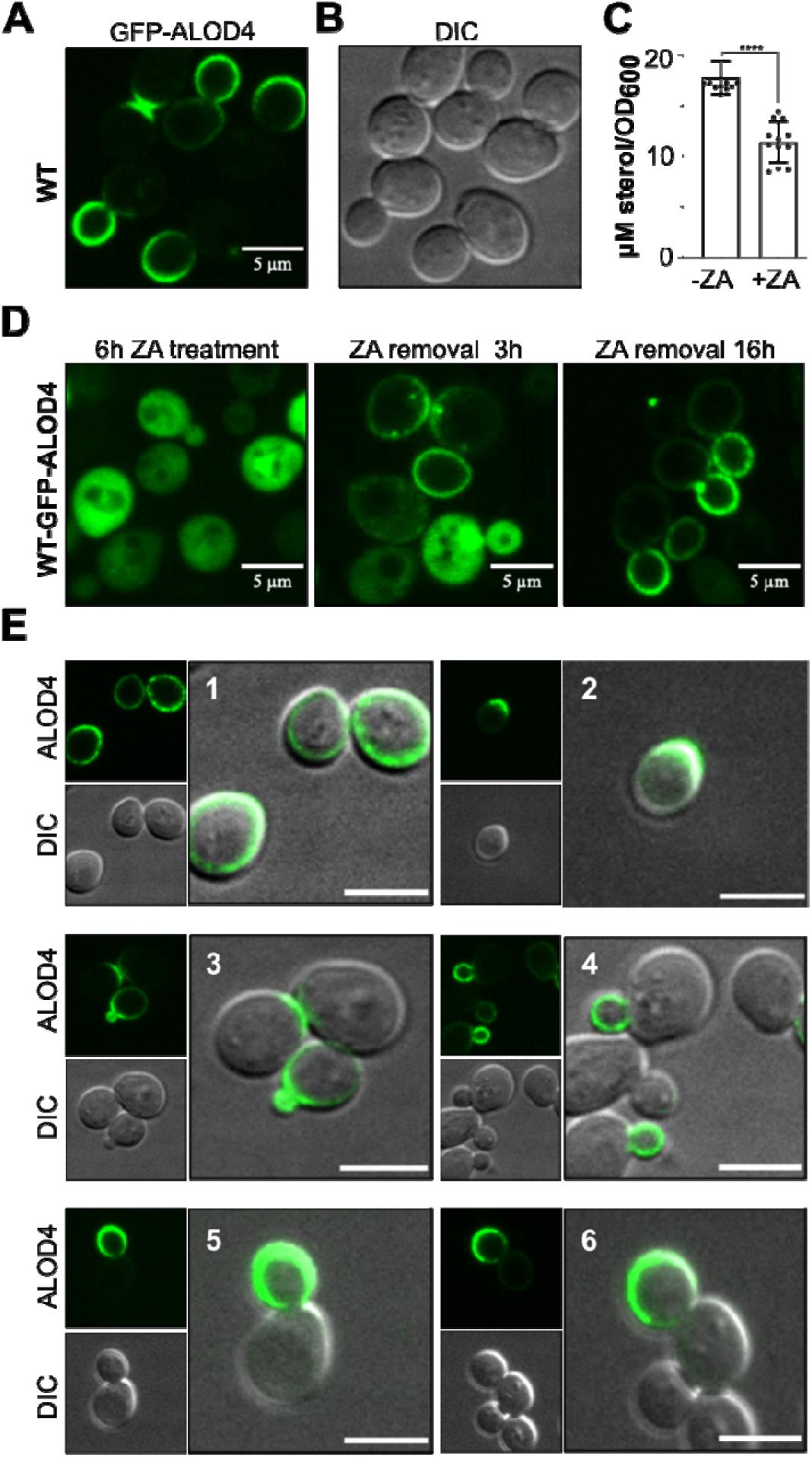
ALOD4 sensor detects sterols in the plasma membrane (PM) and is polarized during cell division. **(A and B)** ALOD4 is a sterol binding sensor that localizes to the PM in logarithmically growing WT cells. Its localization was more prominent in the bud as compared to the mother. **(C)** WT cells were grown in the presence or absence of zaragozic acid for 16h, sterols were extracted and quantified using the amplex red cholesterol oxidase assay (n=3; 12 individual determinations). Sterol levels are reduced by ≈ 38% in the presence of zaragozic acid (P-value <0.0001). **(D)** GFP-ALOD4 localized to the cytosol when WT cells were treated with 10 μg/ml zaragozic acid (ZA) for 6h. Removal of zaragozic acid for the indicated time-points reestablished the PM localization of ALOD4. **(E)** GFP-ALOD4 localized specifically to the bud-tip/bud of the growing bud as the cells progressed through cell division (Panels 1-6). Representative images are shown.

### Ergosterol is polarized during cell division

We observed that the GFP-ALOD4 sensor localizes prominently in the bud compared to the mother cell (Fig.1A, D). In unbudded cells growing in log phase ALOD4 is visible uniformly on the PM (Fig.1E, panel 1). Prior to bud formation the yeast cell becomes polarized and switches to an anisotropic mode of growth at the initiation of cell division. Fig.1E, panel 2 demonstrates the presence of the ALOD4 sensor at the site of initial bud emergence. As the cell moves through cell division (panels 3-6), GFP-ALOD4 localizes predominantly to the growing bud. This data suggests that ergosterol is polarized during cell division in *S. cerevisiae*, revealing a potential role of this central lipid in cell cycle progression. Curiously, filipin does not detect polarization during vegetative growth in *S. cerevisiae* but does reveal inhomogeneous distribution of sterol in *S. pombe* (Takeda et al., 2004; Wachtler et al., 2003). Whether this is due to the sensitivity of filipin, its ability to bind ergosterol in both leaflets of the PM, or that filipin staining is typically done with chemical fixatives that do not immobilize lipids remains unclear.

### Localization of ALOD4 is altered in mutants defective in non-vesicular sterol transport

A recent study has demonstrated that ongoing polarized secretion is required to maintain the PFO**-** D4H on the PM (Encinar Del Dedo et al., 2021). Similar results were previously reported for PtdSer whose polarization required secretion and exclusion of PtdSer from endocytic vesicles (Fairn et al., 2011). We sought to explore the role of non-vesicular sterol transport proteins, Osh and Lam, in the distribution of ALOD4. Since deletion of all seven Osh proteins is incompatible with viability, we used *osh1-7*Δ*(osh4-1*^*ts*^) strain which has a temperature-sensitive mutation in *OSH4*. A shift to a non-permissive temperature of 37°C inactivates the last Osh protein, and as can be seen in Fig.2A, ALOD4 becomes largely cytosolic with a few internal structures also being labeled. In contrast, examination of *ysp2 lam4* cells revealed that while they still have GFP-ALOD4 in the PM, the polarity of the GFP signal is reversed. Previously it was demonstrated that the transport of exogenously supplied sterols in the absence of Lam4 and Ysp2 is reduced by 2-3 fold yet the total ergosterol content of the cell is not affected. However, the localization of our sterol biosensor is drastically altered indicative of its sensitivity to changes in sterol organization of the PM. Since Ysp2 is needed for optimal transport of sterols from the PM to ER and *ysp2* cells are hyper-sensitive to amphotericin B (Gatta et al., 2015; Roelants et al., 2018) the data suggests that in vegetative cells that Ysp2 and Lam4 preferentially remove sterols from the PM of the mother thereby helping to maintain a gradient generated by the secretory pathway.

We sought to determine if other forms of yeast cell polarity also show a gradient in GFP-ALOD4 localization. The formation of mating projections in *S. cerevisiae* is a second well studied form of cell polarity in this organism. Cells expressing RFP-LactC2 and GFP-ALOD4 were treated with mating factor for 2 hours to induce the formation of mating projections and to synchronize the cells cycle. In wildtype cells GFP-ALOD4 was enriched at the tips of mating projections and depleted from the PM of the mother (Fig. 2B). Consistent with the vegetative cells (Fig. 2A) we observed that the *ysp2 lam4* cells have reversed polarity of the ALOD4. Wash out of the mating factor allows cells to enter the cell cycle in a synchronized fashion. Under these circumstances the polarization of the ALOD4 remains reversed in the *ysp2 lam4* cells (Fig. 2B).

**Figure 2.**
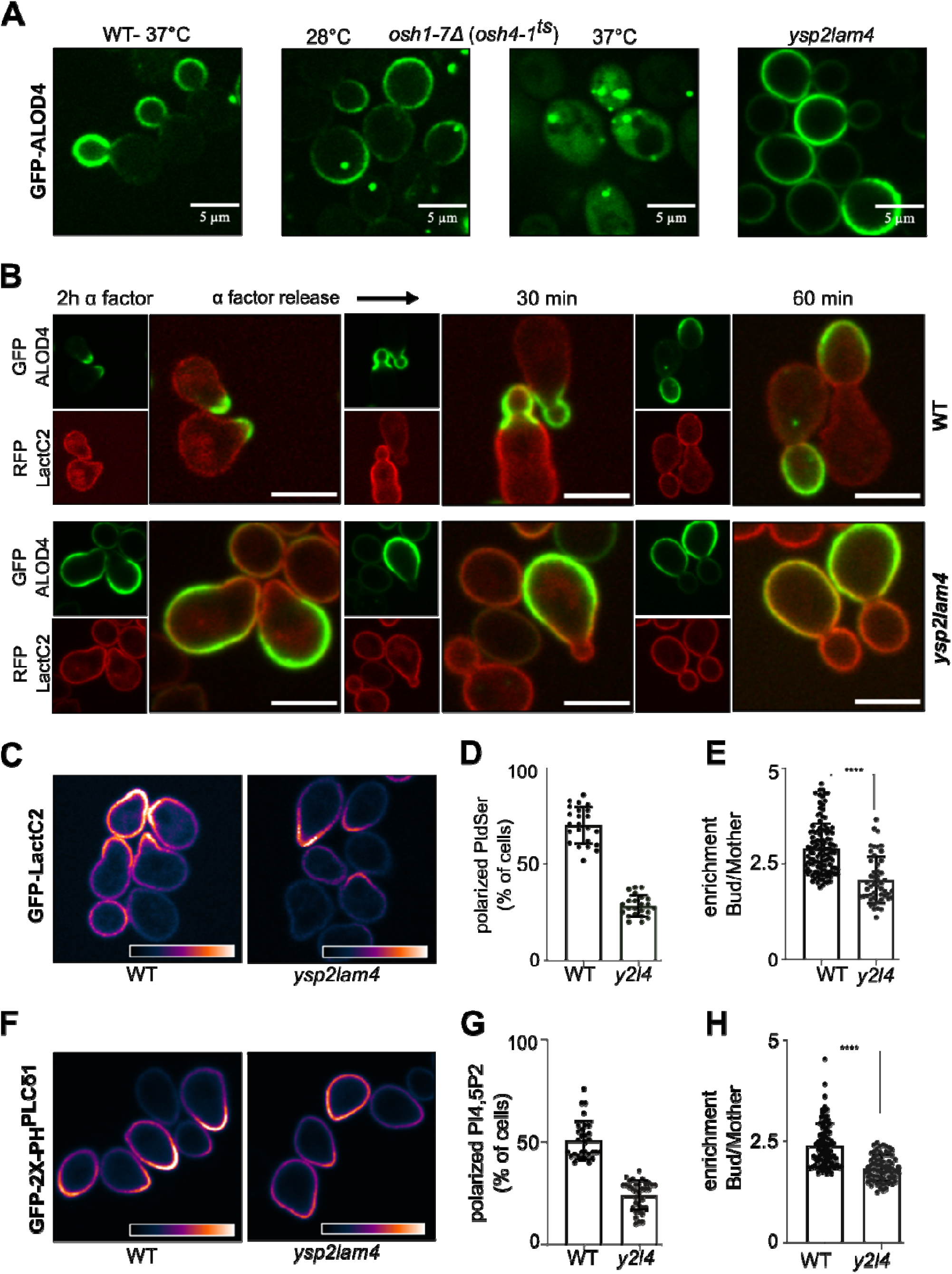
LAM proteins are required for the anisotropic distribution of ALOD4 and optimal polarization of anionic phospholipids. **(A)** Cells deficient in non-vesicular sterol transport - *osh1-7*Δ *(osh4-1*^*ts*^) and *ysp2 lam4* displayed an altered localization of GFP-ALOD4. *osh1-7*Δ*(osh4-1*^*ts*^) mutant at 37ºC showed a cytosolic signal for ALOD4. The *ysp2 lam4* mutant on the other hand showed a signal that was present all over the PM accompanied by a reversal of polarization of ALOD4 compared to WT. **(B)** Polarity of ALOD4 bound sterol is reversed in ysp2 lam4 mutant-WT and *ysp2 lam4* cells were co-labeled with GFP-ALOD4 and RFP-LactC2 and treated with 0.5 μM alpha-factor for 2h to arrest the cell cycle and generate mating projections. The cells were then released into media without alpha-factor, and the cell cycle progression was followed. GFP-ALOD4 localized to the bud-tip after 2h treatment with alpha factor in WT but not in *ysp2 lam4*. From 30-60 min, GFP-ALOD4 was present only in the growing bud in WT; in contrast, GFP-ALOD4 presented only in the mother cell in y*sp2 lam4*. **(C)** WT and *ysp2 lam4* cells were treated with alpha factor for 90 min and the polarization of PtdSer sensor-GFP-LactC2 was analyzed. LUT-gem was applied for visualization of the lipid sensor at the bud-tip. (**D)** The images obtained (in C) were analyzed for the presence of the PtdSer at the site of bud formation; > 500 cells were counted for each cell type. The data is represented as a percentage of polarized cells with polarized PtdSer. **(E)** The fluorescence intensity of the images obtained was determined by measuring the intensity at the polarized tip vs. the non-polarized end of the cell and correcting for the intensity of the cytosolic signal. The data is represented as a ratio of the two intensities. **(F)** WT and *ysp2 lam4* cells were treated with alpha factor for 90 min and the polarization of PI(4,5)P2 sensor-GFP-2×PH(PLCδ1) was analyzed. LUT-gem was applied for visualization of the l PI(4,5)P2 sensor at the bud-tip. **(G)** The images obtained (in F) were analyzed for the presence of the PI(4,5)P_2_ at the site of bud formation; > 500 cells were counted for each cell type. The data is represented as a percentage of polarized cells with polarized PI(4,5)P_2_. **(H)** The fluorescence intensity of the images obtained was determined by measuring the intensity at the polarized tip vs. the non-polarized end of the cell and correcting for the intensity of the cytosolic signal. The data is represented as a ratio of the two intensities. Representative images are shown.

Anionic phospholipids PtdSer, PI4P and PI4,5P_2_ have been reported to be enriched at sites of polarized growth and mating projections where they support the local activation of Cdc42 (Fairn et al., 2011; Garrenton et al., 2010; Meca et al., 2019). We wondered if the decreased sterol abundance at sites of polarized growth, or at least its accessibility to the probe, has a secondary impact on PtdSer and PI4,5P_2_. Wild-type and *ysp2 lam4* cells expressing GFP-LactC2 (PtdSer probe) or GFP-PH-PLCδ (PI4,5P_2_ probe) were treated with mating factor for 90 min and examined by microscopy. Overall, 72% of wildtype and 51% of the *ysp2 lam4* cells displayed shmoos after the 90 min. In wild-type, ∼70% of polarized cells had a >2-fold enrichment of PtdSer at the bud tip compared to ∼28% in *ysp2 lam4* cells. Furthermore, ratiometric analysis of the cells that show a > 2-fold enrichment of PtdSer in the bud/mother revealed that the *ysp2 lam4* cells remained less able to concentrate PtdSer (Fig. 2E). Similarly polarization of PI4,5P_2_ in wild-type was apparent in ∼52% of polarized cells while only ∼24% of the *ysp2 lam4* cells had enrichment of PH-PHCδ (Fig 2F) with even the polarized cells showing less enrichment compared to the wildtype (Fig.2G). The data demonstrate that Ysp2 and Lam4 play a key role in maintaining plasmalemmal lipid gradients during polarized growth.

### The activation of the pheromone response pathway is reduced in the absence of Ysp2 and Lam4

We hypothesized that that disruption of the local lipid environment in the *ysp2 lam4* cells may impair pheromone response and an impairment in mating efficiency. The pheromone response pathway has two signaling modules, a) activation of G protein-coupled receptor pathway and b) the activation MAP kinase cascade. Cdc42 and Ste5 are two downstream targets of the G protein-coupled receptor pathway that are polarized to the bud-tip in the presence of pheromone (Bardwell, 2005). We treated the cells with 0.5 μM α-factor for 2h and investigated the localization of GFP-Cdc42 and GFP-Ste5 in wild-type and *ysp2 lam4* cells. As shown in Fig. 3A, only 37% of polarized *ysp2 lam4* (*y2l4*) cells displayed Cdc42 at the bud-tip compared to wild-type (57%). The polarization of Ste5 was even more reduced with only 17% of *ysp2 lam4* (*y2l4*) cells having Ste5 at the bud-tip compared to wild-type (78%).

**Figure 3.**
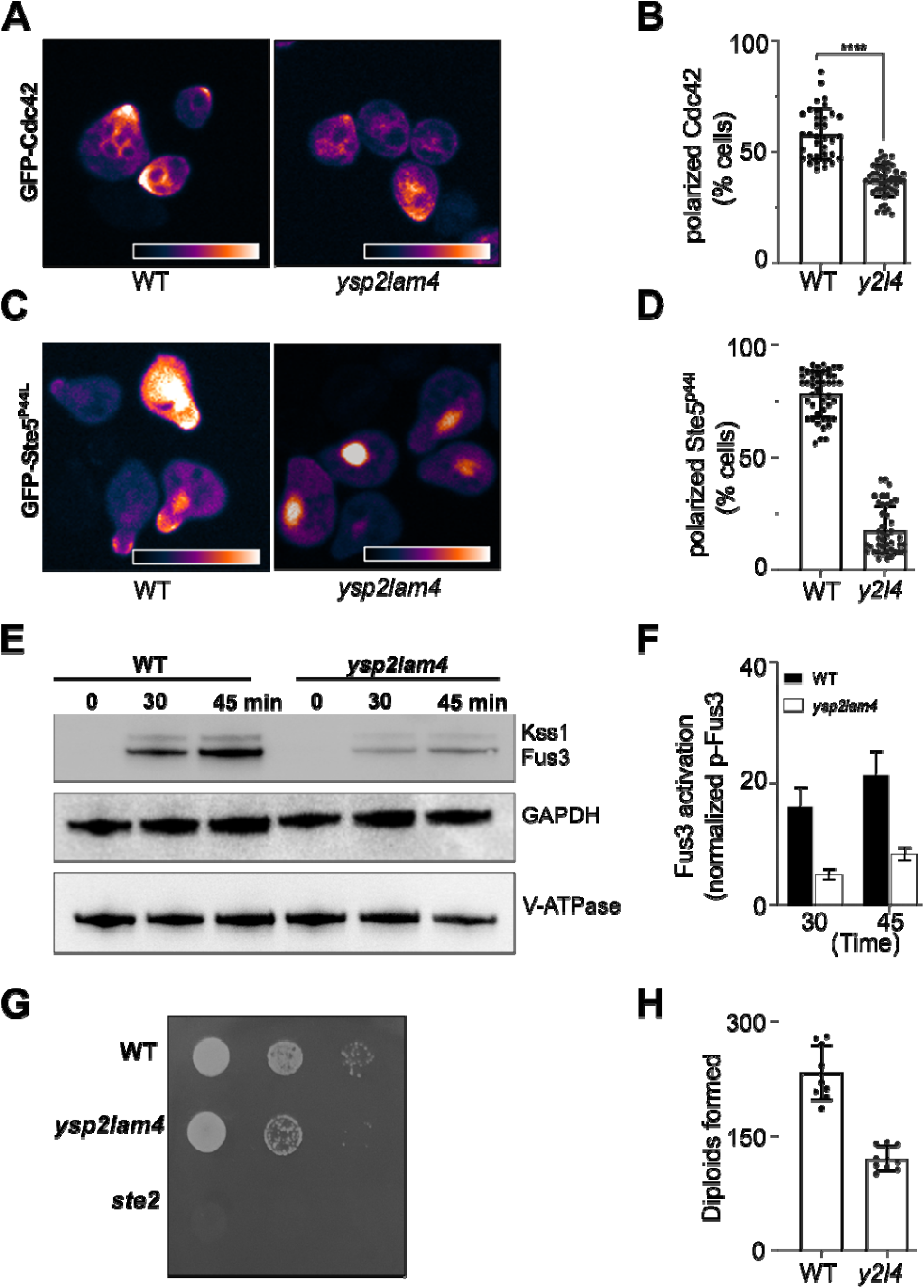
Ysp2 and Lam4 are required for optimal mating and activation of the pheromone response pathway. **(A)** WT and *ysp2 lam4* (*y2l4*) cells expressing GFP-Cdc42 were treated with 0.5 μM alpha-factor for 90 min, and the localization of Cdc42 was analyzed. LUT-gem was applied for visualization of the Cdc42 at the bud-tip. **(B)** The images obtained (in A) were analyzed for the presence of the Cdc42 at the site of bud formation; > 500 cells were counted. The data is represented as a percentage of polarized cells with a polarized Cdc42. GFP-Cdc42 was detected at the bud-tip in 37% of polarized *ysp2 lam4* (*y2l4*) cells compared to 57% in wild-type (P-value <0.0001). **(C)** WT and *ysp2 lam4* (*y2l4*) cells expressing GFP-Ste5^P44L^ were treated with 0.5 μM alpha-factor for 90 min, and the localization of Ste5 was analyzed. LUT-gem was applied for visualization of the Ste5 at the bud-tip. **(D)** The images obtained (in C) were analyzed for the presence of the Ste5 at the site of bud formation; > 500 cells were counted. The data is represented as a percentage of polarized cells with a polarized Ste5. GFP-Ste5 was present at the bud-tip in only 17% of polarized *ysp2 lam4* (*y2l4*) cells compared to 78% in WT (P-value <0.0001). **(E and F)** Stationary phase cells were treated with alpha-factor for 90 min and cell cycle progression was followed for the indicated time-points in the absence of alpha-factor. 5 OD_600_ unit cells were collected at each time point, total protein was extracted, and immunoprecipitation was performed to detect the levels of phosphorylated Fus3 in WT and *ysp2 lam4* mutant. GAPDH and V-ATPase were used as loading controls. The signal intensity of the Fus3 was analyzed using ImageJ (n=3) to determine the extent of Fus3 activation. **(G)** MATa strains of WT and *ysp2 lam4* were incubated with same OD_600_ units of MATα mating tester strain for 4h. For the qualitative mating assay, serial 10-fold dilutions were spotted onto agar plates lacking amino acids such that only diploids would grow. **(H)** To quantify the number diploids formed, WT and *ysp2 lam4* (*y2l4*) strains were incubated with same OD_600_ units of MATα mating tester strain for 4h; approximately 300 cells were spread on amino acid lacking plates and the number of colonies formed for each strain were counted. *ysp2 lam4* mating efficiency was reduced by 48% (n=3).

Ste5 serves as a scaffold protein to allow for a robust MAP kinase signaling cascade. Two MAP kinases activated in response to pheromone are Fus3 and Kss1, with the former being more prominent. To study this module of the pheromone response pathway, we treated stationary phase cells with α-factor for 2h and collected cells for immunoblotting at the indicated times after being released into media lacking the pheromone. As demonstrated in Fig. 3C, the amount of phosphorylated MAP kinase (Fus3) was reduced by 65% in the *ysp2 lam4* cells compared to wildtype. Thus, alterations in the local lipid environment and Ste5 recruitment results in impaired MAP kinase activation in response to mating pheromone.

Since the pheromone response pathway is affected, we hypothesized that the mating efficiency of the *ysp2 lam4* mutant would also be reduced. Fig. 3D, E demonstrates that *ysp2 lam4* has fewer diploids than wild-type, and the mating efficiency is reduced by ∼50%. These data establish that the two sterol transport proteins play critical roles in establishing an ideal lipid environment to promote cell polarity and response to external stimuli.

### Reduced sterol transport in *ysp2 lam4* mutant is responsible for the polarization defects displayed during cell cycle progression

Given the altered localization of the sterol sensor ALOD4 in *ysp2 lam4* cells we postulated that it was due to the reduced of sterol transport function of these proteins. Ysp2 and Lam4 are membrane proteins located at ER-PM contact sites and contain two sterol-binding StARkin domains reported to play a role in sterol transport and homeostasis (Gatta et al., 2015; Jentsch et al., 2018). The second more StARkin domain (Lam4S2, Fig. 4A) has been previously studied and transports dehydroergosterol *in vitro* at a rate of ∼0.84 molecules / molecule of Lam4S2 per second (Jentsch et al., 2018). Expression of this domain in isolation is sufficient to rescue the nystatin/amphotericin sensitivity of the *ysp2 lam4* mutant(Khelashvili et al., 2019). Elucidation of its crystal structure revealed residue K89, K1031 in the full-length protein, to be critical for sterol transport (Jentsch et al., 2018). Mutation of this residue to alanine (K89A) not only abolishes in vitro sterol transport but also fails to correct the nystatin sensitivity (Khelashvili et al., 2019).

**Figure 4.**
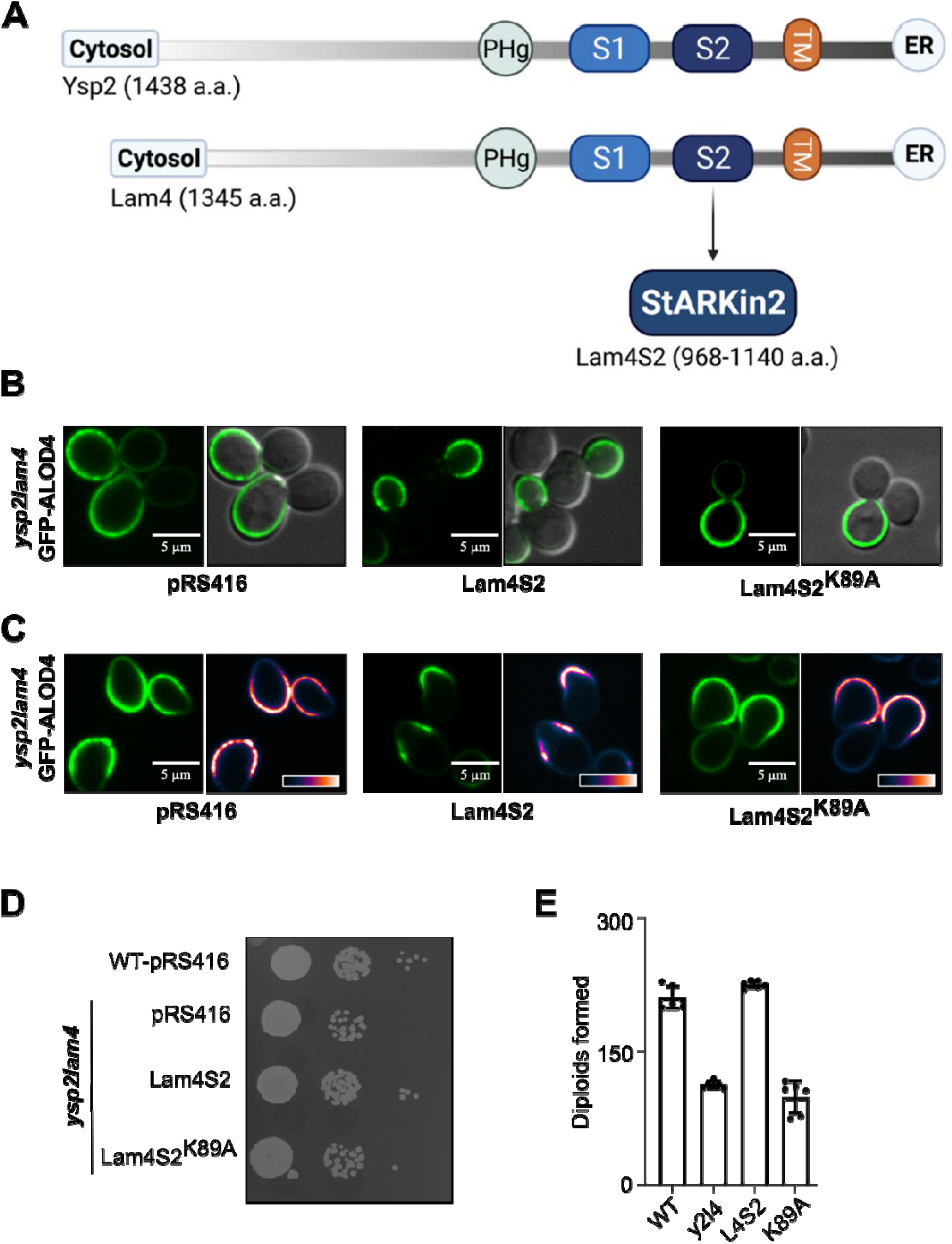
The second StARkin domain of Lam4 rescues the polarization defect of ysp2lam4 mutant. **(A)** Ysp2 and Lam4 are anchored via their transmembrane domain (TM) to endoplasmic reticulum (ER) at sites where ER is in close contact with the PM. They have a PH (pleckstrin homology)-like GRAM domain (PHg) and two StART-like StARKin domains (S1 and S2). The function of StARKin2 domain of Lam4 (Lam4S2) has been extensively studied. **(B and C)** *ysp2 lam4* mutant was transformed with GFP-ALOD4 and an empty vector (pRS416) or with a vector for the expression of Lam4S2 wild-type or point mutant K89A. In the presence of a functional StARkin domain, the polarity defect of the mutant is rescued (Lam4S2). The defect persists when the domain is non-functional (Lam4S2^K89A^). This is observed in both logarithmically growing cells (B) and cells treated with alpha-factor for 90 min (C). **(D)** MATa strains of WT containing the empty plasmid and *ysp2lam4* expressing indicated plasmids were incubated with same OD_600_ units of MATα mating tester strain for 4h. For the qualitative mating assay serial 10-fold dilutions were spotted onto agar plates lacking amino acids such that only diploids would grow. **(E)** To quantify the number of diploids, WT (pRS416), *ysp2 lam4*+pRS416 (*y2l4*), *ysp2 lam4*+Lam4S2 (L4S2) and *ysp2 lam4+*Lam4S2^K89A^ (K89A) strains were incubated with same OD_600_ units of MATα mating tester strain for 4h; approximately 300 cells were spread on amino acid lacking plates and the number of colonies formed for each strain were counted. Lam4S2 recovers the mating efficiency defect of *ysp2 lam4* (n=3).

To determine if the reversal of sterol polarization in *ysp2 lam4* mutant was explicitly due to its reduced sterol transport function, we introduced the functional StARkin domain (Lam4S2) and a non-functional version (Lam4S2^K89A^) into *ysp2 lam4* and analyzed the localization of GFP-ALOD4. In *ysp2 lam4* cells that were in the logarithmic phase (Fig. 4B) or treated with α-factor to induce polarization (Fig. 4C), GFP-ALOD4 localization to the bud/mating project was corrected in the presence of Lam4S2. In contrast, in the presence of Lam4S2^K89A^, GFP-ALOD4 failed to accumulate at the sites of polarized growth (Fig. 4B). In addition to the ability of the Lam4S2 to re-establish the enrichment of sterols to the tips of the mating projects, the mating efficiency defect of *ysp2 lam4* was also recovered in the presence of Lam4S2 but not Lam4S2^K89A^ (Fig. 4D and E).

Ysp2 and Lam4 are required to maintain the lipid homeostasis of the PM (Gatta et al., 2015). They transport sterols in vitro (Jentsch et al., 2018; Khelashvili et al., 2019) and their depletion at contact sites in tether mutants results in reduced levels of critical PM lipids (Quon et al., 2018). The results suggest that the ability of LAM proteins to transfer sterols from the PM to the ER may support sterol anisotropy in two ways. First, selective removal of ergosterol from the mother would help maintain a front-to-back polarization. Second, returning ergosterol to the ER for re-entry into the secretory pathway will aid in its delivery to sites of polarized growth and mating. Considering that their sterol transport function is attenuated via TORC2 mediated phosphorylation (Roelants et al., 2018), another possibility is that phosphorylation of Ysp2 and Lam4 occur preferentially in the bud preventing the removal of ergosterol from the sites of polarized growth. These findings establish the role of GRAM proteins in sterol transport and in the polarization of sterols during cell cycle progression. Our data further reinstates this; they are required for establishing polarity-dependent gradients of ergosterol, PS, and PIP2-all critical lipids of the PM.

## Materials and methods

### Strains and Growth Conditions

The *Saccharomyces cerevisiae* strains used in this study are listed in Table 1. Yeast cells were grown in standard rich (YPD) and defined minimal (SC) media containing 2% glucose and supplemented with appropriate nutrients to maintain selection for plasmids. All cells expect *osh1-7* Δ (*osh4-1*^*ts*^) were grown at 30ºC which was moved to 37ºC for 1h for inactivation of Osh4 prior to microscopy. Plasmids pRS416-GFP-Lam4S2 and pRS416-GFP-Lam4S2^K89A^ was a gift from Anant Menon (Khelashvili et al., 2019). The GFP encoded by these plasmids was mutated G67A to make a non-fluorescent variant (Bartkiewicz et al., 2018).

### Microscopy

Imaging in the St. Michael’s Hospital BioImaging centre was conducted using spinning-disc confocal systems (Quorum Technologies). An Axiovert 200M microscope (Carl Zeiss) with ×63 objective lenses and a ×1.5 magnifying lens was equipped with diode-pumped solid-state lasers (440, 491, 561, 638, and 655 nm; Spectral Applied Research) and a motorized XY stage (Applied Scientific Instrumentation). Images were acquired using back-thinned, electron-multiplied cameras (model C9100-13 ImagEM; Hamamatsu Photonics). Images obtained were analyzed using Fiji-ImageJ (Schindelin et al., 2012).

### Inhibitor treatment

α-Mating Factor acetate salt and zaragozic acid were obtained from Sigma Aldrich and Cayman chemical, respectively. Cells were treated with 10 μg/ml (w/v) zaragozic acid for 6 -16 h followed by microscopy or lipid extraction. To study cell cycle progression, cells were treated with 0.5 μM α-factor (prepared in DMSO) for 90-120 min as indicated. Following the treatment, α-factor was removed, and cells were allowed to proceed through the cell cycle for 60 min. Cells were harvested every 30 min and immediately visualized by microscopy. To analyze the polarization of lipid probes (GFP-LactC2 and GFP-2X-PH-PLCδ1), GFP-Cdc42, and GFP-Ste5P44L (a gift from Peter Pryciak (Winters et al., 2005)) cells were treated with 0.6 μM α-factor for 90 min and visualized by microscopy. GFP-Ste5^P44L^ was induced by moving the cells into media containing 2% galactose 30 min prior to adding alpha-factor to the media. The images obtained were analyzed using ImageJ.

### Lipid extraction and Amplex Red assay for determination of cellular sterol content

10 OD_600_ units of wild-type yeast cells grown either in the presence or absence of 10 μg/ml zaragozic acid for 16 h were harvested and the lipids were extracted from the intact yeast cells as previously described (Chauhan et al., 2019). The extracted lipids were resuspended in 500 μl methanol; 5 μl of the lipid extract was used to estimate the total sterol content using the Amplex™ Red Cholesterol Assay Kit (Invitrogen) per the manufacturer’s instructions.

### Preparation of Yeast Whole-Cell Extracts and Immunoblotting

Whole cell extracts were prepared as previously described by (Baerends et al., 2000; Chauhan et al., 2015). Briefly, cells were synchronized by treating with 0.5 μM α-factor for 2h and released into fresh media (no α-factor). 5 OD_600_ units of cells were harvested at the indicated time points, collected by centrifugation, resuspended in 400 μl of 12.5% (w/v) trichloroacetic acid (TCA) and incubated at –80 °C overnight to allow protein precipitation. TCA was removed by centrifugation, and the cell pellets were washed twice with 80% (v/v) ice-cold acetone, air-dried, and dissolved in 1% SDS/0.1N NaOH and 1× SDS loading buffer, samples were boiled for 5 min and proteins were resolved on 10% SDS polyacrylamide gels and blotted onto nitrocellulose membranes (Bio-Rad). Anti-rabbit Phospho-p44/42 MAPK antibody (Cell Signaling Technology) was used to detect phosphorylated Fus3 and Kss1. Since antibody for detection of total Fus3/Kss1 is not available; anti-GAPDH (Proteintech) and anti-V-ATPase 60 kDa subunit (Invitrogen) were used as loading control. HRP conjugated anti-rabbit IgG and anti-mouse IgG (Cell Signaling Technology) were the secondary antibodies, and ECL Western Blotting Substrate (Pierce) was used for detection. The signal intensity of the immunoblots was calculated using ImageJ.

### Mating assay

Mat-α tester strain and the strains to be tested for their mating efficiency were grown overnight in appropriate media. 1 OD_600_ unit of each strain was mixed with the same OD_600_ unit of the mat-α tester strain in 200 μl YPD and incubated at room temperature without shaking for 4h to allow mating. 10-fold serial dilutions of the mated strains were spotted onto agar plates lacking amino acids and incubated at 28ºC for 48h. To determine the mating efficiency, approximately 300-400 cells from the second serial dilution were added to 50 μl autoclaved water; spread plated on amino acid lacking agar plates and grown at 28ºC for 48h. The number of colonies formed were counted.

## Supplemental Material

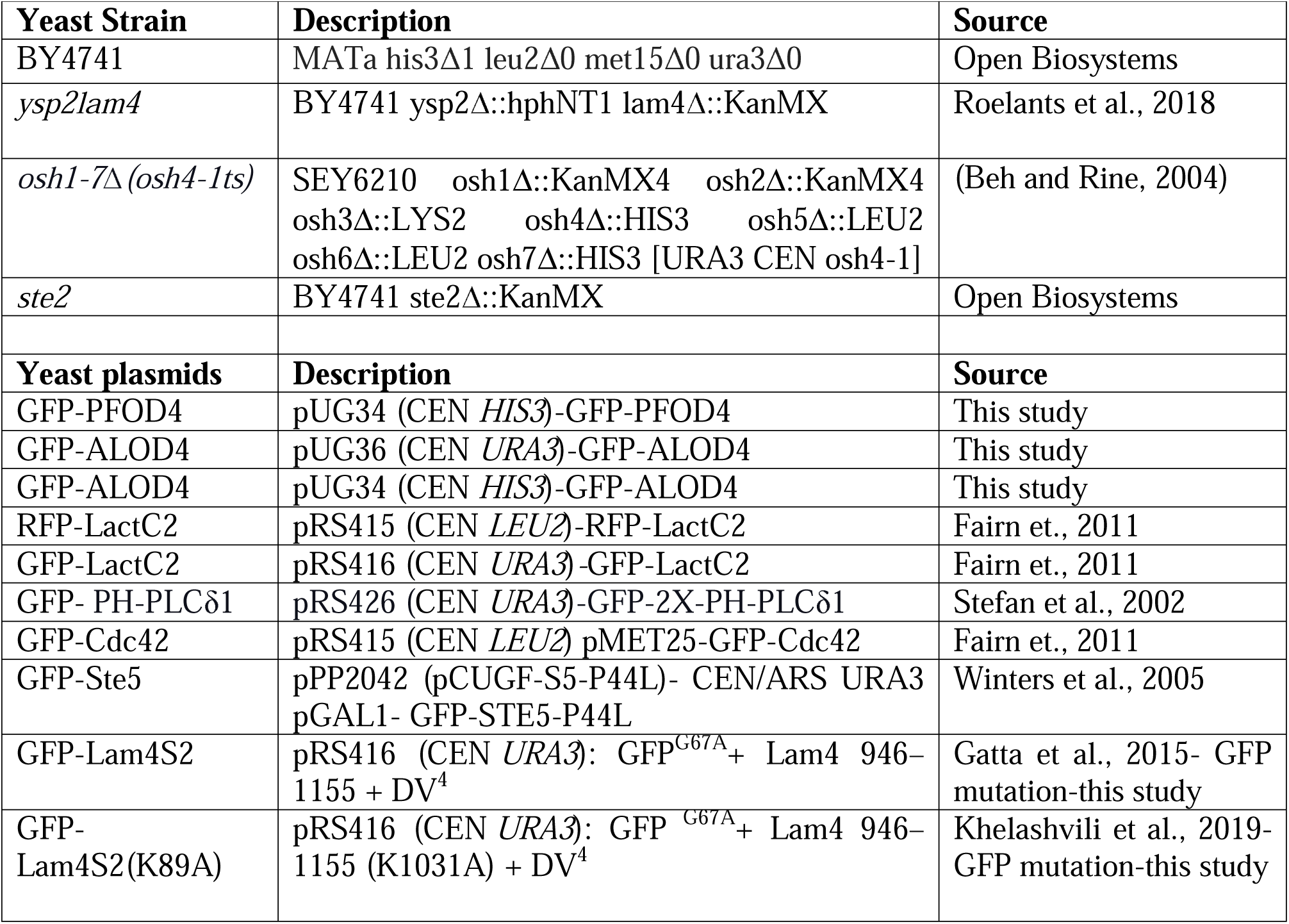

## Acknowledgments

We thank Dr. Peter Pryciak (University of Massachusetts Medical School) and Dr. Anant K. Menon (Weill Cornell Medicine) for the plasmids. The authors would like to acknowledge the Kennan Research Center for Biomedical Science Core Facilities at St. Michael’s Hospital, especially Dr. Caterina Di Ciano-Oliveira for their continued technical advice, expertise, and training. **Funding:** This work was supported by an NSERC Discovery Grant RPGIN-2019-04425 to GDF. We also would like to thank Ms. Hannah Wallworth for assistance analyzing the ratiometric images. Acknowledge Hannah for image analysis

## References

Baerends, R.J., K.N. Faber, A.M. Kram, J.A. Kiel, I.J. van der Klei, and M. Veenhuis. 2000. A stretch of positively charged amino acids at the N terminus of Hansenula polymorpha Pexp is involved in incorporation of the protein into the peroxisomal membrane. J Biol Chem. 275:9986–9995.

Bagnat, M., and K. Simons. 2002. Cell surface polarization during yeast mating. Proc Natl Acad Sci U S A. 99:14183–14188.

Bardwell, L. 2005. A walk-through of the yeast mating pheromone response pathway. Peptides. 26:339–350.

Bartkiewicz, M., S. Kazazic, J. Krasowska, P.L. Clark, B. Wielgus-Kutrowska, and A. Bzowska. 2018. Non-fluorescent mutant of green fluorescent protein sheds light on the mechanism of chromophore formation. FEBS Lett. 592:1516–1523.

Beh, C.T., L. Cool, J. Phillips, and J. Rine. 2001. Overlapping functions of the yeast oxysterol-binding protein homologues. Genetics. 157:1117–1140.

Chauhan, N., J.A. Jentsch, and A.K. Menon. 2019. Measurement of Intracellular Sterol Transport in Yeast. Methods Mol Biol. 1949:115–136.

Chauhan, N., M. Visram, A. Cristobal-Sarramian, F. Sarkleti, and S.D. Kohlwein. 2015. Morphogenesis checkpoint kinase Swe1 is the executor of lipolysis-dependent cell-cycle progression. Proc Natl Acad Sci U S A. 112:E1077–1085.

Encinar Del Dedo, J., I.M. Fernandez-Golbano, L. Pastor, P. Meler, C. Ferrer-Orta, E. Rebollo, and M.I. Geli. 2021. Coupled sterol synthesis and transport machineries at ER-endocytic contact sites. J Cell Biol. 220.

Fairn, G.D., M. Hermansson, P. Somerharju, and S. Grinstein. 2011. Phosphatidylserine is polarized and required for proper Cdc42 localization and for development of cell polarity. Nat Cell Biol. 13:1424–1430.

Flanagan, J.J., R.K. Tweten, A.E. Johnson, and A.P. Heuck. 2009. Cholesterol exposure at the membrane surface is necessary and sufficient to trigger perfringolysin O binding. Biochemistry. 48:3977–3987.

Garrenton, L.S., C.J. Stefan, M.A. McMurray, S.D. Emr, and J. Thorner. 2010. Pheromone-induced anisotropy in yeast plasma membrane phosphatidylinositol-4,5-bisphosphate distribution is required for MAPK signaling. Proc Natl Acad Sci U S A. 107:11805–11810.

Gatta, A.T., L.H. Wong, Y.Y. Sere, D.M. Calderon-Norena, S. Cockcroft, A.K. Menon, and T.P. Levine. 2015. A new family of StART domain proteins at membrane contact sites has a role in ER-PM sterol transport. Elife. 4.

Hodge, R.G., and A.J. Ridley. 2016. Regulating Rho GTPases and their regulators. Nat Rev Mol Cell Biol. 17:496–510.

Infante, R.E., and A. Radhakrishnan. 2017. Continuous transport of a small fraction of plasma membrane cholesterol to endoplasmic reticulum regulates total cellular cholesterol. Elife. 6.

Jentsch, J.A., I. Kiburu, K. Pandey, M. Timme, T. Ramlall, B. Levkau, J. Wu, D. Eliezer, O. Boudker, and A.K. Menon. 2018. Structural basis of sterol binding and transport by a yeast StARkin domain. J Biol Chem. 293:5522–5531.

Jin, H., J.M. McCaffery, and E. Grote. 2008. Ergosterol promotes pheromone signaling and plasma membrane fusion in mating yeast. J Cell Biol. 180:813–826.

Johnson, B.B., P.C. Moe, D. Wang, K. Rossi, B.L. Trigatti, and A.P. Heuck. 2012. Modifications in perfringolysin O domain 4 alter the cholesterol concentration threshold required for binding. Biochemistry. 51:3373–3382.

Khelashvili, G., N. Chauhan, K. Pandey, D. Eliezer, and A.K. Menon. 2019. Exchange of water for sterol underlies sterol egress from a StARkin domain. Elife. 8.

Kozubowski, L., K. Saito, J.M. Johnson, A.S. Howell, T.R. Zyla, and D.J. Lew. 2008. Symmetry-breaking polarization driven by a Cdc42p GEF-PAK complex. Curr Biol. 18:1719–1726.

Maekawa, M., and G.D. Fairn. 2015. Complementary probes reveal that phosphatidylserine is required for the proper transbilayer distribution of cholesterol. J Cell Sci. 128:1422–1433.

Marek, M., V. Vincenzetti, and S.G. Martin. 2020. Sterol biosensor reveals LAM-family Ltc1-dependent sterol flow to endosomes upon Arp2/3 inhibition. J Cell Biol. 219.

Meca, J., A. Massoni-Laporte, D. Martinez, E. Sartorel, A. Loquet, B. Habenstein, and D. McCusker. 2019. Avidity-driven polarity establishment via multivalent lipid-GTPase module interactions. EMBO J. 38.

Orlando, K., and W. Guo. 2009. Membrane organization and dynamics in cell polarity. Cold Spring Harb Perspect Biol. 1:a001321.

Quon, E., Y.Y. Sere, N. Chauhan, J. Johansen, D.P. Sullivan, J.S. Dittman, W.J. Rice, R.B. Chan, G. Di Paolo, C.T. Beh, and A.K. Menon. 2018. Endoplasmic reticulum-plasma membrane contact sites integrate sterol and phospholipid regulation. PLoS Biol. 16:e2003864.

Roelants, F.M., N. Chauhan, A. Muir, J.C. Davis, A.K. Menon, T.P. Levine, and J. Thorner. 2018. TOR complex 2-regulated protein kinase Ypk1 controls sterol distribution by inhibiting StARkin domain-containing proteins located at plasma membrane-endoplasmic reticulum contact sites. Mol Biol Cell. 29:2128–2136.

Schindelin, J., I. Arganda-Carreras, E. Frise, V. Kaynig, M. Longair, T. Pietzsch, S. Preibisch, C. Rueden, S. Saalfeld, B. Schmid, J.Y. Tinevez, D.J. White, V. Hartenstein, K. Eliceiri, P. Tomancak, and A. Cardona. 2012. Fiji: an open-source platform for biological-image analysis. Nat Methods. 9:676–682.

Slaughter, B.D., S.E. Smith, and R. Li. 2009. Symmetry breaking in the life cycle of the budding yeast. Cold Spring Harb Perspect Biol. 1:a003384.

Takeda, T., T. Kawate, and F. Chang. 2004. Organization of a sterol-rich membrane domain by cdc15p during cytokinesis in fission yeast. Nat Cell Biol. 6:1142–1144.

Wachtler, V., S. Rajagopalan, and M.K. Balasubramanian. 2003. Sterol-rich plasma membrane domains in the fission yeast Schizosaccharomyces pombe. J Cell Sci. 116:867–874.

Wedlich-Soldner, R., S.C. Wai, T. Schmidt, and R. Li. 2004. Robust cell polarity is a dynamic state established by coupling transport and GTPase signaling. J Cell Biol. 166:889–900.

Winters, M.J., R.E. Lamson, H. Nakanishi, A.M. Neiman, and P.M. Pryciak. 2005. A membrane binding domain in the ste5 scaffold synergizes with gbetagamma binding to control localization and signaling in pheromone response. Mol Cell. 20:21–32.

Witte, K., D. Strickland, and M. Glotzer. 2017. Cell cycle entry triggers a switch between two modes of Cdc42 activation during yeast polarization. Elife. 6.

Woods, B., C.C. Kuo, C.F. Wu, T.R. Zyla, and D.J. Lew. 2015. Polarity establishment requires localized activation of Cdc42. J Cell Biol. 211:19–26.

